# Genetic Characterization of Glyoxalase Pathway in Oral Streptococci and its Contribution to Interbacterial Competition

**DOI:** 10.1101/2023.08.07.552317

**Authors:** Lin Zeng, Payam Noeparvar, Robert A Burne, Benjamin S. Glezer

**Affiliations:** Department of Oral Biology, University of Florida College of Dentistry, Gainesville, Florida, USA

**Keywords:** Reactive Electrophile Species, Methylglyoxal, Dental Caries, Streptococcus mutans, PTS, competition

## Abstract

Substantial quantities of Reactive Electrophile Species (RES), including methylglyoxal and glyoxal, are generated by microbes and humans. To understand the impact of RES on oral microbial homeostasis, genetic analyses were performed on the glyoxalase pathway in *Streptococcus mutans* (SMU) and *Streptococcus sanguinis* (SSA). Loss of glyoxalase I (LguL), which catalyzes the rate-limiting reaction in RES degradation, reduced methylglyoxal and glyoxal tolerance to a far greater extent in SMU than in SSA, decreasing the competitiveness of SMU over SSA in planktonic cultures. MICs showed an overall greater RES tolerance by SMU than SSA; a finding consistent with the ability of methylglyoxal to induce the expression of *lguL* in SMU, but not in SSA. Computational analysis identified a novel paralogue of LguL in most streptococci represented by SMU.1112c in SMU. ΔSMU.1112c showed a minor decrease in methylglyoxal tolerance under certain conditions, but a significant growth defect on fructose; a phenotype reversed by the deletion of a fructose-1-phosphate-generating sugar: phosphotransferase system or addition of glutathione (GSH) to the medium. Further, deletion of the glucose-PTS in SMU increased RES tolerance partly through enhanced expression of the pyruvate-dehydrogenase complex. Consistent with the requirement of GSH for methylglyoxal detoxification, deletion of glutathione synthetase (*gshAB*) in SMU significantly reduced RES resistance. This study reveals the critical roles of RES in fitness and interbacterial competition and the effects of PTS in modulating RES metabolism. The fact that RES may impact the pathogenic potential of the oral microbiome via differential effects on beneficial and pathogenic species warrants further investigation.

**Importance:** As highly reactive byproducts of sugar metabolism, very little is known regarding the contribution of methylglyoxal or related aldehyde compounds to oral health. The need to better understand the influence of these reactive electrophile species (RES) to microbial physiology and ecology is made more urgent by the widespread condition of hyperglycemia in humans, which is associated with elevated RES levels. Our study showed a significantly greater ability of a major caries pathobiont, *Streptococcus mutans*, to tolerate methylglyoxal and glyoxal than many commensal oral streptococci. Genetic analysis of methylglyoxal degradation in the pathobiont and commensals identified significant differences in genetic structure and gene regulation patterns that could contribute to differential fitness by constituents of the dental microbiota and ecologic shift in the presence of RES.

## Introduction

The initial pathway for catabolism of carbohydrates for energy production is the Embden-Meyerhof-Parnas pathway, often referred to as glycolysis. Incoming sugars are phosphorylated and, by the activities of a series of enzymes conserved in both bacteria and mammalian cells, converted into pyruvate and a cohort of metabolic intermediates that are critical to bacterial growth and metabolism (1). Either catalyzed by specific enzymes or produced as the byproducts of various non-enzymatic reactions, reactive electrophile species (RES), including two prominent dicarbonyl compounds, methylglyoxal (MG) and glyoxal (GO), are created during metabolism of sugars, proteins or lipids (2). These RES metabolites are often produced in significant quantities in cells actively engaged in carbohydrate metabolism, or in diabetic patients experiencing poor metabolic/glycemic control (3). Due to their biochemical properties, RES are membrane-permeable and highly reactive towards most cellular macromolecules, including proteins, nucleic acids, and lipids (4, 5). These reactions can significantly alter the nature and functions of affected biological molecules, and often result in the creation of other reactive intermediates and advanced glycation end-products (AGEs); both RES and AGEs are implicated in the pathophysiology of chronic conditions such as aging, diabetes, and cancer development (6). For example, MG and GO contribute to the progression of type 2 diabetes (T2DM) and development of its many complications, such as kidney diseases (diabetic nephropathy), vision (retinopathy) and neurological damages (3). Because of their reactivity, the majority of RES are present in bound forms to biological molecules, so their abundance is frequently underestimated. Using a highly sensitive approach that targets MG in both free form and bound to biological ligands, it was recently determined that in a mammalian cell line total MG levels can be as high as 310 μM, whereas previous estimations were 1-10 μM in various biological systems (7). Similarly high levels of MG have been reported after treatment of the periodontal pathogen *Tannerella forsythia* with glucose, or during infection of macrophages by certain mycobacteria (8, 9).

While most bacteria have the necessary apparatus to detoxify and degrade MG and GO, some (e.g., *Escherichia coli*, *Bacillus subtilis*) actively engage in biosynthesis of MG as a mechanism, termed methylglyoxal bypass, to control glycolytic rates, especially under high-sugar or low-phosphate conditions (4, 10, 11). In this process, a metabolic intermediate of glycolysis, dihydroxyacetone phosphate (DHAP) is dephosphorylated and converted into MG, which upon a spontaneous reaction with the reduced form of glutathione (GSH), forms a hemithioacetal that is then converted into S-lactoylglutathione (SLG) by the activity of glyoxalase I. SLG is subsequently converted into D-lactate and GSH via the activity of glyoxalase II. D-lactate can then be oxidized by a unique D-lactate dehydrogenase to pyruvate to end the MG bypass. SLG can also serve as an important allosteric effector that activates a potassium: proton antiporter KefGB of *E. coli*, which in turn acidifies the cytoplasm and enhances RES tolerance (12). However, most Gram-positive species lack the enzyme required for biosynthesis of MG, methylglyoxal synthase (MgsA) (13, 14). Instead, glyoxalases I and II in these bacteria work in concert to degrade MG originating exogenously or created by endogenous metabolic activities, during which the activity of glyoxalase I is often the rate-limiting step (12, 15, 16). Furthermore, because of the need for GSH during the metabolism of MG, exposure to MG can disrupt the redox homeostasis of GSH and GSSG (17).

Recent advances have highlighted the complex interplay between human microbiomes and systemic health. With T2DM affecting close to 10% of the world’s adult population (18), there is an urgent need for better understanding of the influence of the pathophysiological state of T2DM on the genomics, biochemistry, and ecology of the human microbiome. For example, studies of periodontal diseases in the context of diabetes have established a two-way relationship between T2DM and microbiome dysbiosis that is often the driving force behind deteriorating periodontal health (19). For dental caries, there has been an increasing body of evidence associating poor glycemic control with increased risk for dental caries (20–25). As such, more in-depth research is needed to address the impact of increased excretion of RES on microbial homeostasis in the context of dental caries. As the most abundant members of the supragingival microbiota and potent producers of organic acids, streptococci contribute directly to dental health and diseases by shaping the taxonomic and biochemical landscape of the biofilm (26–28). In this report, we performed genetic analysis on several genes whose products are essential to RES tolerance of a model caries pathobiont, *Streptococcus mutans*. Our study revealed the ability of RES to influence streptococcal competition, as well as the interconnected nature of PTS and RES metabolism.

## Results and Discussion

### Identification of putative MG metabolic genes in streptococci

Apart from biochemical studies in the model microorganisms *E. coli* and *B. subtilis*, relatively little is known regarding the contribution and regulation of MG metabolism in microbial pathogenesis (3). To begin to better understand MG metabolism in the oral microbiome, we conducted a search for putative MG synthetic and metabolic genes in several important streptococci, including *S. mutans* (SMU), *Streptococcus sanguinis* (SSA), *Streptococcus gordonii* (SGO), *Streptococcus mitis,* and *Streptococcus pneumoniae*, etc. As depicted in Fig. 1, in addition to genes predicted to encode for glyoxalase I (*lguL/gloA1*) and glyoxalase II (*gloB*), another glyoxalase I paralogue (tentatively named *gloA2*) was identified in most Gram-positive species analyzed other than *S. mitis* and *S*. *pneumoniae*. *gloA2* orthologues were also identified in isolates of *E. coli* as an uncharacterized gene. Aside from *Enterococcus faecalis*, most lactic acid bacteria do not appear to harbor a gene for MG synthase (*mgsA*).

**Fig. 1.**
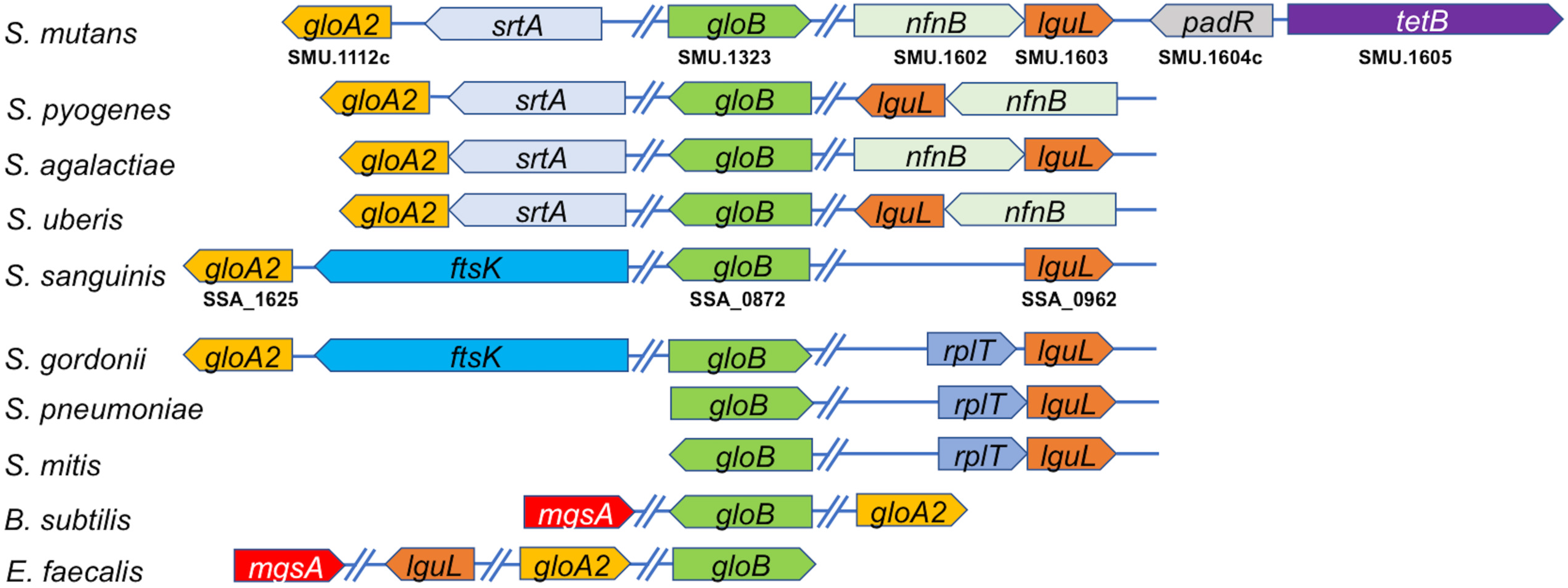
Genetic organization of genes/ORFs likely involved in MG metabolism in important Gram-positive bacteria. Bacteria depend on the glyoxalase pathway for metabolism of MG and GO, a two-step reaction that is catalyzed by gene products of *lguL* (*gloA1,* SMU.1603) and *gloB* (SMU.1323). Based on sequence and structure similarity (Fig. 2), a paralogue of *lguL* (*gloA2,* SMU.1112c) with no known function has been identified in most species analyzed.

The crystal structure of SMU.1112c/GloA2 from *S. mutans* has been solved (Protein Data Bank, RCSB PDB). It is a Zn^2+^-binding protein predicted to be active as a homodimer. SMU.1112c and the predicted structure of SMU.1603/LguL/GloA1 from the same bacterium are 29% identical in sequence and show high degrees of structural similarity (Fig. 2 and Fig. S1), supporting that both proteins may function in RES metabolism. In *S. mutans* and several other streptococci, SMU.1603/*lguL* forms an apparent operon structure with another ORF encoding a putative NAD(P)H-flavin oxidoreductase (SMU.1602)(Fig. 1), a conclusion supported by our RT-qPCR (below) and multiple publications of transcriptomic and proteomic analyses that showed co-regulation of these two genes under various conditions (29, 30). Downstream and in opposite orientation to that of SMU.1602-1603 is a putative transcription regulator, SMU.1604c that shares homology with the PadR family repressors (31). Adjacent to SMU.1604c and in opposing orientation is SMU.1605, a putative transmembrane protein sharing homology with a drug efflux pump of *Bacillus*. Notably different from SMU.1602-SMU.1604c in *S. mutans*, no transcription regulator gene was identified in similar locations in other streptococci. Incidentally, SMU.1604c was also crystalized and determined to be an Mg^2+^-binding protein (RCSB PDB).

**Fig. 2.**
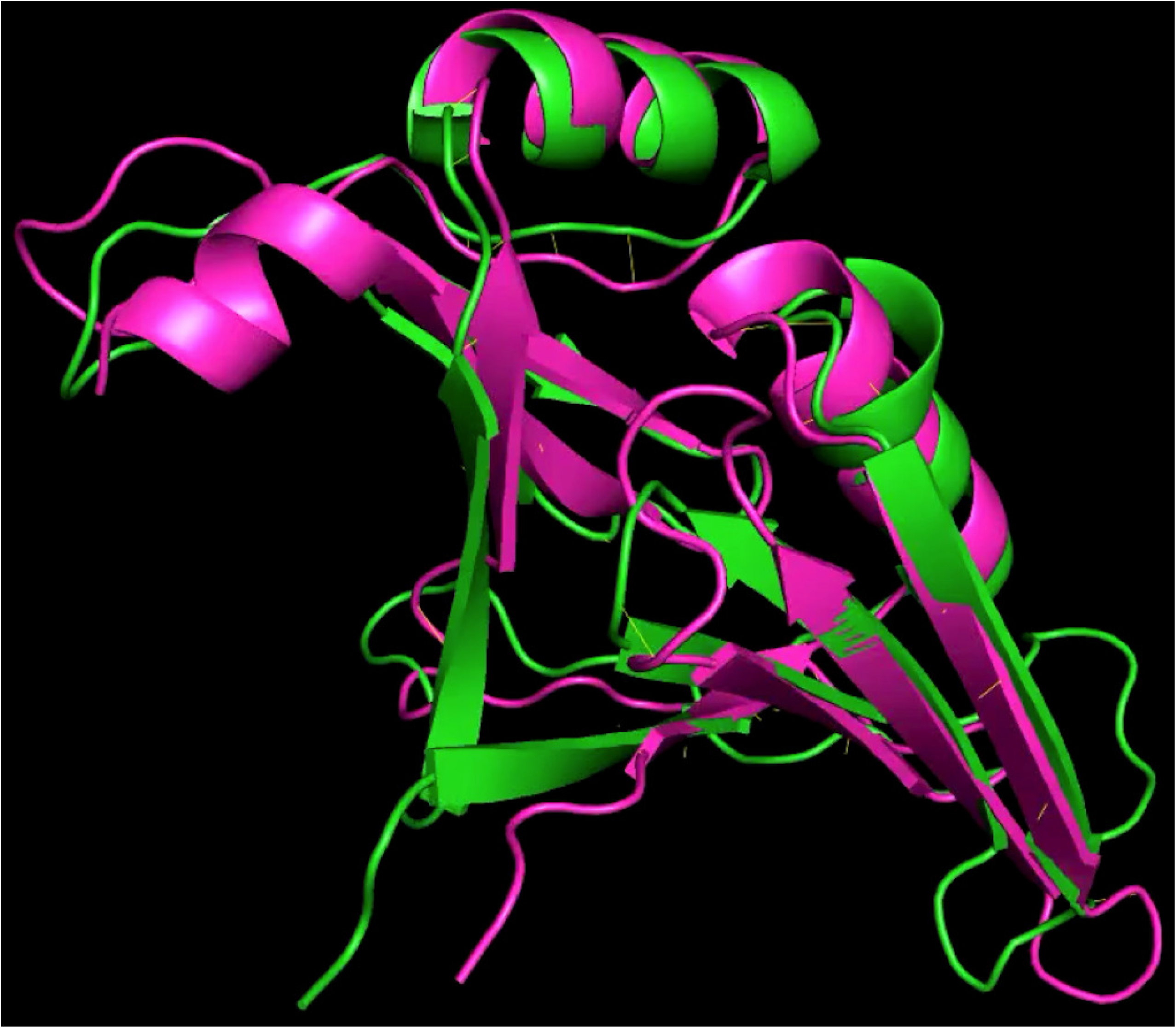
Structure overlay between SMU.1112c (green) and SMU.1603 (purple). The structure of SMU.1603 was rendered using Alphafold 2.

### Genetic analyses in *S. mutans* and *S. sanguinis* of genes required for MG metabolism

To assess the genetic structure and function of each gene/ORF predicted to degrade MG, individual deletions were engineered in *S. mutans* strain UA159 (SMU UA159) by knocking out SMU.1112c (*gloA2*), SMU.1603 (*lguL*), and SMU.1323 (*gloB*), followed by phenotypic characterization associated with RES tolerance. At the same time, deletion of equivalent genes, SSA_0962 (*lguL*) and SSA_1625 (*gloA2*), were constructed similarly in the background of *S. sanguinis* SK36. These deletion mutants and their knock-in complementation derivatives were analyzed in planktonic growth assays utilizing a synthetic medium (FMC) supplemented with MG or GO, and MICs for MG or GO were determined by a micro-dilution assay in FMC.

Relative to the wild type SMU UA159, deletion of *lguL* (SMU.1603) resulted in a reduction in tolerance to MG (Table 1) from 5.5 mM to 1.5 mM (∼75%), and an extended lag phase in FMC-glucose supplemented with 1 mM of MG (Fig. 3). These results confirmed previous studies (13, 14) that suggested that LguL is the major GloA enzyme responsible for degrading MG in streptococci. However, different from a previous report on LguL in *S. mutans* (14), ΔSMU.1603 did not show any significant change in decrease in its ability to grow in FMC-glucose adjusted to pH 5.5 or 6.0, nor did any other mutants analyzed here (Fig. S2). Also consistent with previous findings in other bacterial systems (15), loss of SMU.1323, encoding a putative GloB enzyme, resulted in no reduction in MG tolerance (data not shown).

**Fig. 3.**
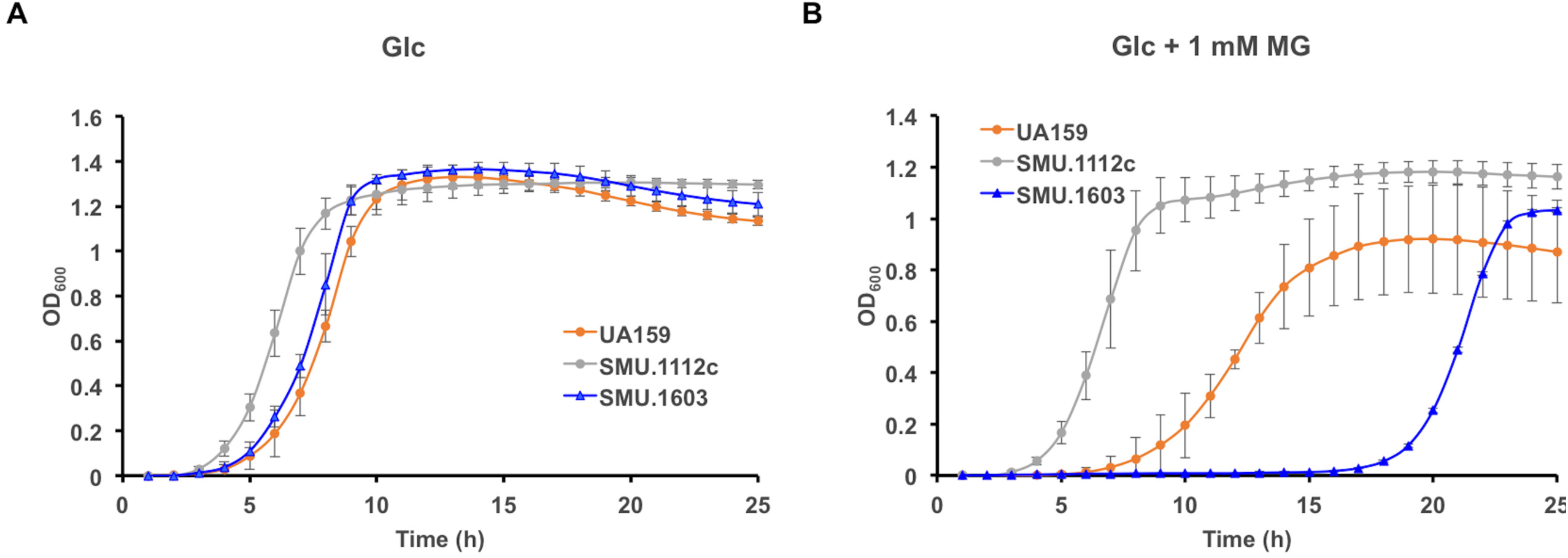
Growth curves of SMU strain UA159 and its isogenic mutants deficient in SMU.1112c or SMU.1603. Bacterial cultures prepared with BHI were diluted into a synthetic FMC medium constituted with 20 mM of glucose and 0 (A) or 1 mM (B) of MG. Each strain was represented by at least three individual cultures. All cultures were covered with sterile mineral oil and incubated at 37°C for monitoring of optical density at 600 nm (OD_600_).

**Table 1.**
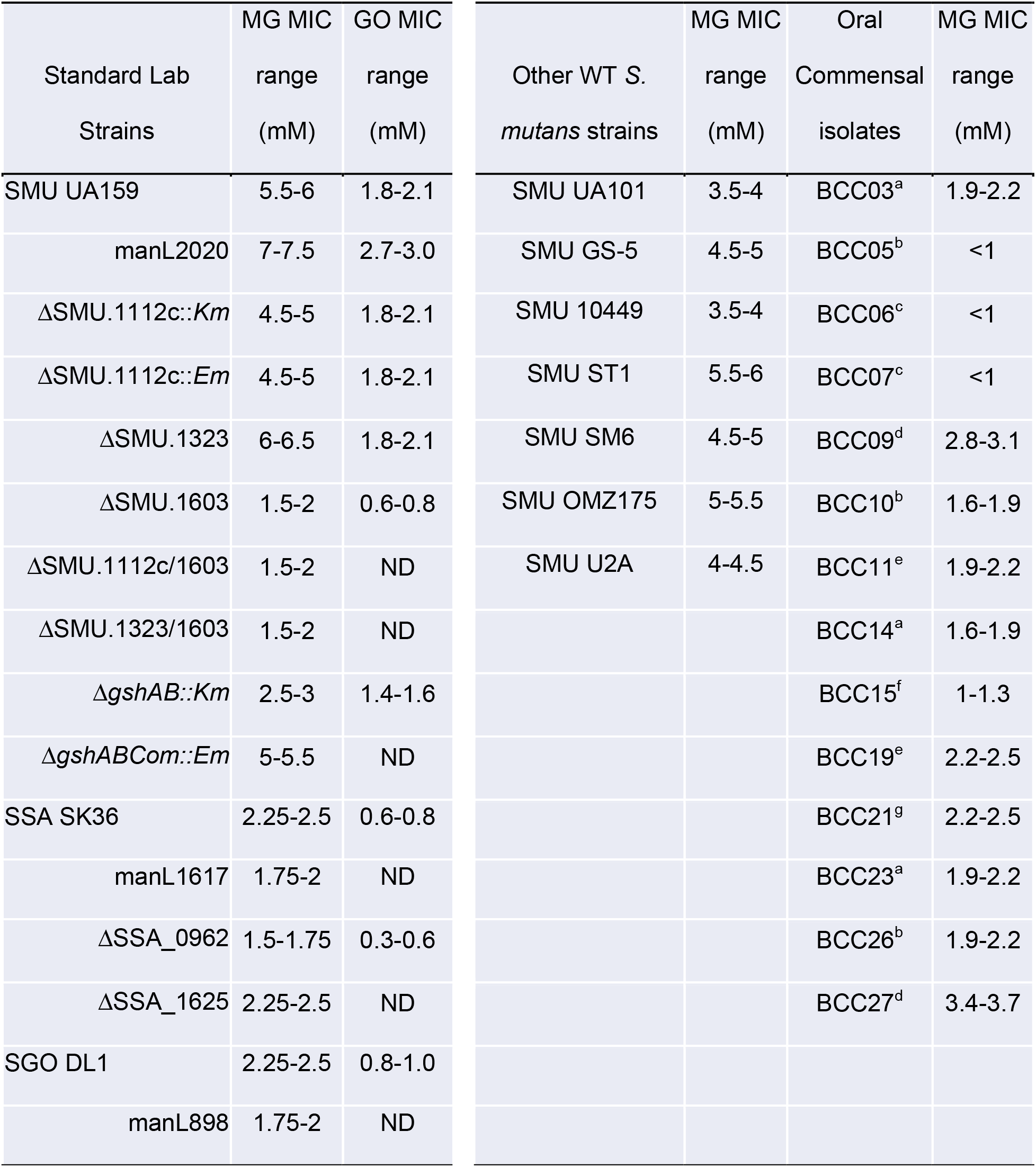
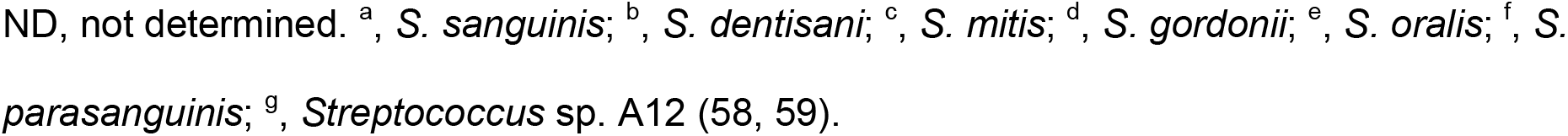
MIC (minimum inhibitory concentrations) of MG and GO measured in wild-type oral streptococcal strains and mutant derivatives. Each result is presented as a range to denote both the average MIC (first number) and the next increment of concentration.

| Standard Lab<br>Strains | MG MIC<br>range<br>(mM) | GO MIC<br>range<br>(mM) | Other WT S.<br>*mutans* strains | MG MIC<br>range<br>(mM) | Oral<br>Commensal<br>isolates | MG MIC<br>range<br>(mM) |
| --- | --- | --- | --- | --- | --- | --- |
| SMU UA159 | 5.5-6 | 1.8-2.1 | SMU UA101 | 3.5-4 | BCC03<sup>a</sup> | 1.9-2.2 |
| manL2020 | 7-7.5 | 2.7-3.0 | SMU GS-5 | 4.5-5 | BCC05<sup>b</sup> | <1 |
| ΔSMU.1112c::*Km* | 4.5-5 | 1.8-2.1 | SMU 10449 | 3.5-4 | BCC06<sup>c</sup> | <1 |
| ΔSMU.1112c::*Em* | 4.5-5 | 1.8-2.1 | SMU ST1 | 5.5-6 | BCC07<sup>c</sup> | <1 |
| ΔSMU.1323 | 6-6.5 | 1.8-2.1 | SMU SM6 | 4.5-5 | BCC09<sup>d</sup> | 2.8-3.1 |
| ΔSMU.1603 | 1.5-2 | 0.6-0.8 | SMU OMZ175 | 5-5.5 | BCC10<sup>b</sup> | 1.6-1.9 |
| ΔSMU.1112c/1603 | 1.5-2 | ND | SMU U2A | 4-4.5 | BCC11<sup>e</sup> | 1.9-2.2 |
| ΔSMU.1323/1603 | 1.5-2 | ND | | | BCC14<sup>a</sup> | 1.6-1.9 |
| ΔgshAB::*Km* | 2.5-3 | 1.4-1.6 | | | BCC15<sup>f</sup> | 1-1.3 |
| ΔgshABCom::*Em* | 5-5.5 | ND | | | BCC19<sup>e</sup> | 2.2-2.5 |
| SSA SK36 | 2.25-2.5 | 0.6-0.8 |  |  | BCC21<sup>g</sup> | 2.2-2.5 |
| manL1617 | 1.75-2 | ND |  |  | BCC23<sup>a</sup> | 1.9-2.2 |
| ΔSSA_0962 | 1.5-1.75 | 0.3-0.6 | | | BCC26<sup>b</sup> | 1.9-2.2 |
| ΔSSA_1625 | 2.25-2.5 | ND | | | BCC27<sup>d</sup> | 3.4-3.7 |
| SGO DL1 | 2.25-2.5 | 0.8-1.0 |  |  |  |  |
| manL898 | 1.75-2 | ND |  |  |  |  |

ND, not determined. <sup>a</sup>, *S. sanguinis*; <sup>b</sup>, *S. dentisani*; <sup>c</sup>, *S. mitis*; <sup>d</sup>, *S. gordonii*; <sup>e</sup>, *S. oralis*; <sup>f</sup>, *S.* *parasanguinis*; <sup>g</sup>, *Streptococcus* sp. A12 (58, 59).

Deletion of SMU.1112c slightly reduced the MIC for MG (from 5.5 mM to 4.5 mM). Deletion of SMU.1112c or SMU.1323 in the background of ΔSMU.1603 did not further increase its susceptibility to MG. Surprisingly, strain ΔSMU.1112c grew significantly better in planktonic growth curve assays than the wild type did on FMC-glucose containing 1 mM MG; with a shorter lag phase, faster growth rate and higher final optical density (Fig. 3). ΔSMU.1112c also grew slightly better than the wild type on glucose without addition of MG. It appears that the functionality of SMU.1112c in MG resistance is significantly influenced by the condition under which the assays are conducted. Notably, MIC assays were performed using static cultures maintained in ambient atmosphere supplemented with 5% CO_2_, whereas the cultures for growth curve assays were covered by mineral oil and no additional CO_2_ in the atmosphere.

Interestingly, two important commensal streptococci, *S. sanguinis* and *S. gordonii*, were significantly less tolerant to treatment by MG (Table 1, Fig. S3). Similar to UA159, deletion of SSA_0962 (*lguL*) in the background of SSA SK36 resulted in lower MIC values for MG (Table 1) when growing in FMC-glucose. Notably different from SMU was the fact that deletion of SSA_0962 resulted in less than a 1-mM reduction in MIC for MG, as opposed to a 4-mM drop caused by deletion of SMU.1603 in SMU background, although strains ΔSSA_0962 and ΔSMU.1603 now have similar tolerance against MG. Deletion of SSA_1625, the orthologue of SMU.1112c, failed to show any change in MG tolerance. We further expanded the MIC assay to include 7 additional wild-type SMU isolates and 14 clinical isolates of various species of commensal streptococci. The results (Table 1) showed that, compared to most commensal streptococcal species tested, each *S. mutans* strain was significantly more tolerant to MG. Last, likely more reactive than MG by having two aldehyde groups, glyoxal (GO) exerted greater growth inhibition than MG when used at the same levels (Table 1). Again SMU UA159 was notably more tolerant to GO than SSA SK36 or SGO DL1, and genetic deletions that altered MG tolerance in both species similarly impacted relative tolerance to GO, a finding consistent with their chemical similarity and likely shared pathways that bacteria use for detoxification (2). Thus, there appears to be a significant difference in the contributions of *lguL* gene products to MG tolerance among oral streptococci.

### SMU.1112c and PTS are involved in RES metabolism in a metabolite-specific manner

To better understand the phenotypes of strain ΔSMU.1112c, further analyses were carried out by growing it on different carbohydrates. When observed under the microscope, ΔSMU.1112c cells presented significantly longer chain length than the wild type, formed clusters/aggregates, and tended to fall out of the culture media (Fig. 4A). This phenotype was apparent in liquid BHI medium and FMC-fructose, but not in FMC containing other tested sugars (Fig. S4). As cell chaining and clustering is often associated with stress, we performed another growth assay using fructose as the supporting sugar. As shown in Fig. 5AB, ΔSMU.1112c had a slower growth rate and lower final yield than the wild type, as well as lower CFU counts from the same cultures. Measurements of extracellular DNA (eDNA) in the culture supernates of ΔSMU.1112c also showed significantly higher levels of eDNA than the wild type after 24-h in FMC-fructose, consistent with increased cell lysis (Fig. 5C). Also, development of biofilms by SMU UA159 was negatively affected by the deletion of SMU.1112c in a 48-h biofilm assay using BM medium containing 2 mM sucrose and 18 mM glucose (Fig. 5D). ΔSMU.1112c also produced less biofilm on BM containing 2 mM sucrose and 18 mM fructose (Fig. S5). Interestingly, deletion of SMU.1323 and SMU.1603 similarly resulted in lower biofilm mass on BM containing glucose. To contrast the phenotype of ΔSMU.1112c with wild-type cells under RES stress, we treated exponentially growing SMU UA159 cells with 1 mM MG. After 1 h of incubation, cells treated with MG showed a slight but consistent increase in chain length under the microscope (Fig. 4B).

**Fig. 4.**
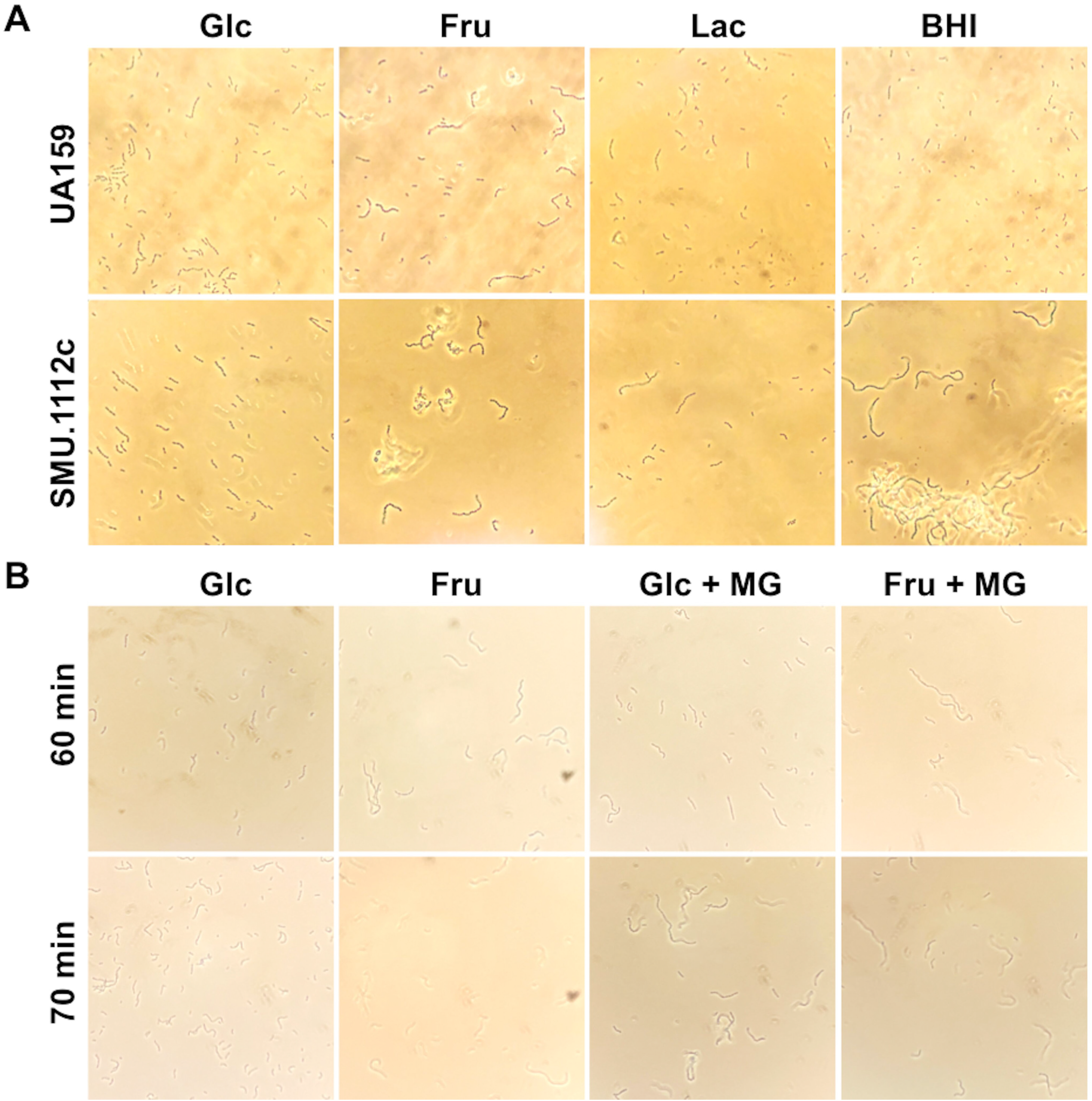
Phase-contrast microscopy. (A) Strains UA159 and ΔSMU.1112c were each cultured in BHI, or FMC with glucose (Glc), fructose (Fru), or lactose (Lac) overnight. (B) Exponential-phase cultures of UA159 (OD_600_ = 0.4) prepared in FMC with Glc or Fru were treated with 1 mM MG for 60 or 70 min.

**Fig. 5.**
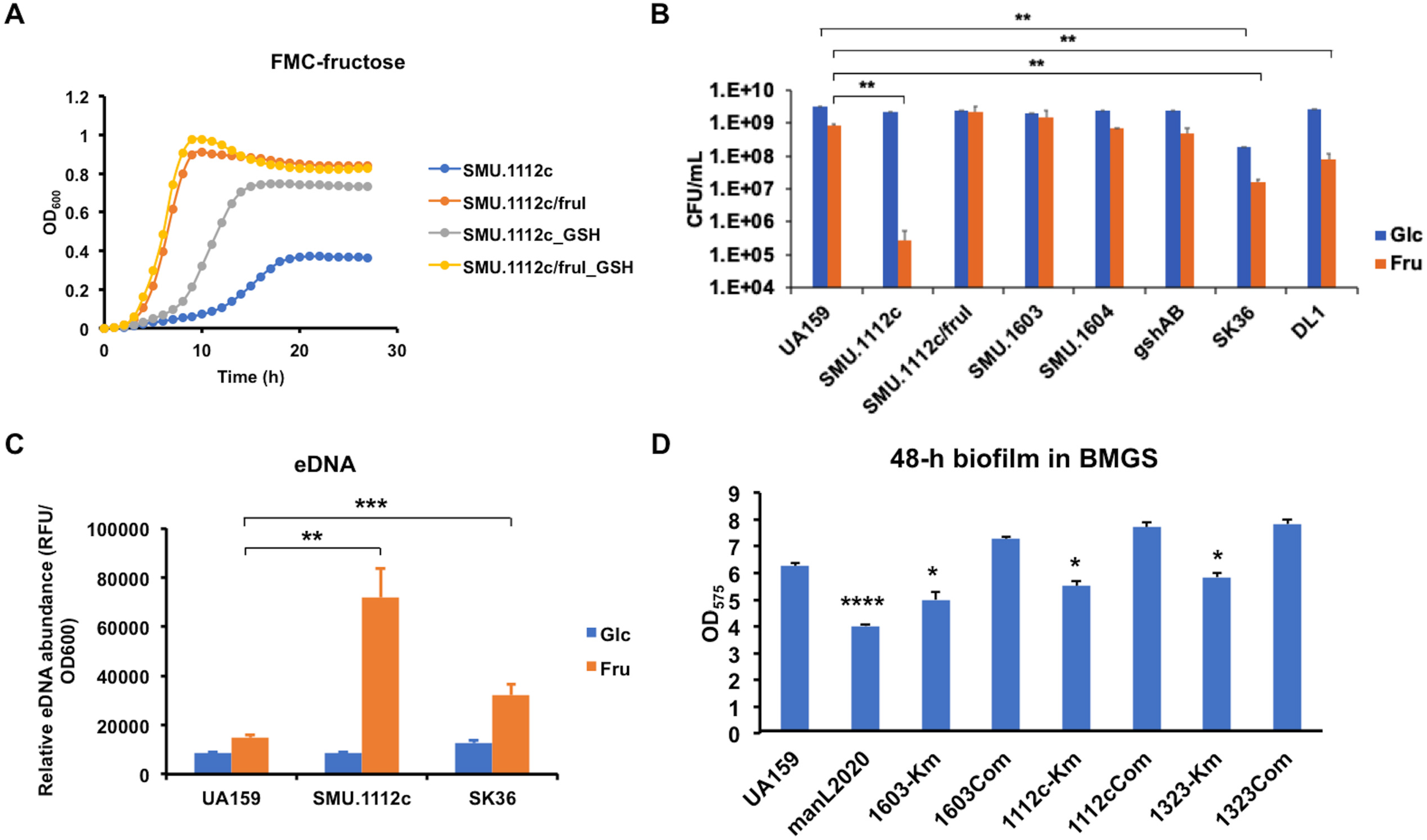
Phenotypic characterization of mutants deficient in methylglyoxal and related metabolic genes. (A) Growth curves of strains ΔSMU.1112c and ΔSMU.1112c/*fruI* in FMC supported with 20 mM fructose, with or without 1 mM GSH. (B) CFU counts of 24-h bacterial cultures in FMC supported with 20 mM glucose or fructose. (C) Relative eDNA abundance in 24-h cultures in FMC supported with glucose or fructose, normalized against the final OD_600_. (D) Total biomass of 48-h biofilms measured as optical density (OD_575_) of crystal violet stain. Each sample was represented by at least three independent cultures. Asterisks denote statistical significance assessed by Student’s *t*-test (*, *P* < 0.05; **, *P* < 0.01; ***, *P* < 0.001; ****, *P* < 0.0001).

Our previous research on fructose metabolism by *S. mutans* (32, 33) showed the significance of fructose-1-phosphate (F-1-P) as a potential stress signal that led to enhanced autolysis and release of DNA, although the mechanism remains to be clarified. To test the theory that the growth phenotype of strain ΔSMU.1112c is influenced by F-1-P accumulation, we deleted the PTS EII permease, FruI that is responsible for generating most of F-1-P in *S. mutans* (32) in the SMU.1112c-null background. A mutant lacking another fructose-PTS transporter, LevDEFG (34) which yields F-6-P, in the SMU.1112c-null background was also included for comparison. Loss of *fruI*, but not *levD* (Fig. S6), reversed the growth phenotype of ΔSMU.1112c in fructose-based medium, including the growth rate and yield (Fig. 5A), and extent of chaining (Fig. 6). Regarding the phenotype of ΔSMU.1112c when growing on BMGS and BHI, it has been demonstrated that sucrose catabolism by *S. mutans* often leads to release of free fructose; BHI medium contains a small amount of fructose contaminant (35, 36).

**Fig. 6.**
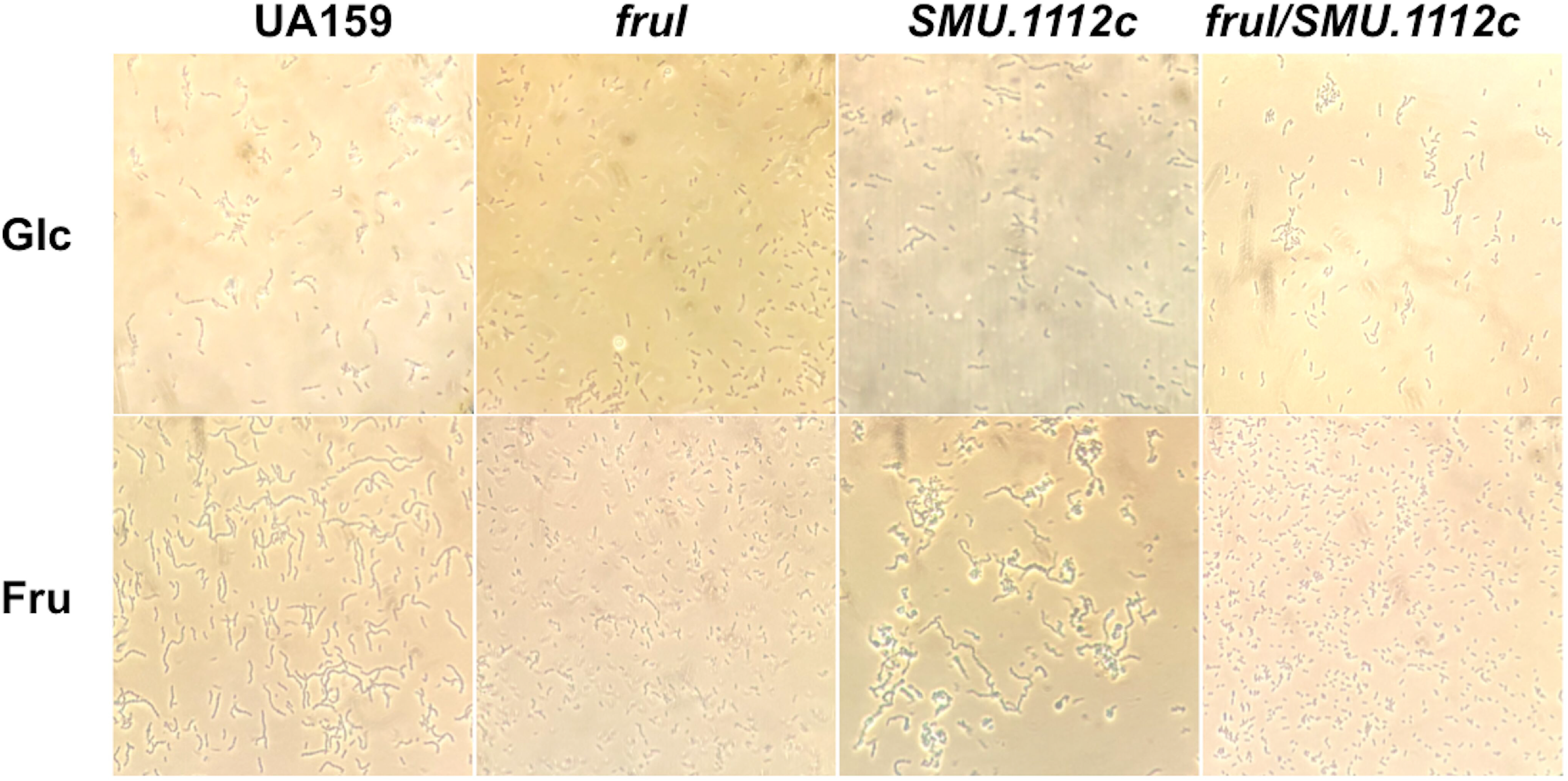
Loss of a F-1-P-generating PTS (FruI) reversed the phenotype of ΔSMU.1112c. Overnight cultures of strain ΔSMU.1112c in FMC with 20 mM Glc or Fru were subjected to phase-contrast microscopy.

We next tested the possibility that F-1-P mediated stress phenotype involves RES generation, which requires the activity of SMU.1112c for detoxification. It is understood that fructose, when metabolized as F-1-P, can bypass the allosteric regulation exerted onto the phosphofructokinase (PFK-1) during glycolysis which uses F-6-P as a substrate, resulting in faster generation of RES (37). Further, research has suggested that oxidative degradation of fructose under the influence of Fenton reaction, i.e., hydrogen peroxide (H_2_O_2_) in the presence of Fe^2+^, strongly potentiates the creation of RES compound GO and fructose-mediated cytotoxicity in mammalian cells (38, 39).

Since reduced glutathione (GSH) is the substrate required for detoxification of MG and GO, we added varying amounts of GSH to the cultures of ΔSMU.1112c in FMC-fructose. When observed under the microscope, GSH reversed the clumping phenotype of ΔSMU.1112c in a dose-dependent manner (Fig. 7); addition of GSH also clearly rescued its growth phenotype on fructose (Fig. 5A). Last, we tested a deletion mutant of UA159 deficient in glutathione synthetase (*gshAB*)(40) in its RES tolerance. The result showed a MIC for MG at 2.5-3 mM, a level slightly higher than ΔSMU.1603, but significantly lower than the wild-type parent (Table 1). Therefore, it appears that the fitness phenotype of strain ΔSMU.1112c involves F-1-P and the depletion of cellular GSH pools. However, since strain ΔSMU.1603 does not display a similar phenotype when grown with fructose, nor does strain ΔSMU.1112c display GO sensitivity, the mechanism by which SMU.1112c confers protection against the metabolites of F-1-P likely differs that of glyoxalases I and II as we understand. To explore the role of *gloA2* orthologue in commensal streptococci, deletion mutant of SSA_1625 in SSA was compared to the wild type SSA SK36 for growth on different sugars. Relative to SMU UA159, SSA SK36 released greater amounts of eDNA when growing on fructose and had lower persistence when the CFUs of the cultures were compared (Fig. 5BC). Like its counterpart in SMU background, cells of strain ΔSSA_1625 fell out of solution in BHI cultures and showed longer chain length than the wild type SK36 (Fig. S7).

**Fig. 7.**
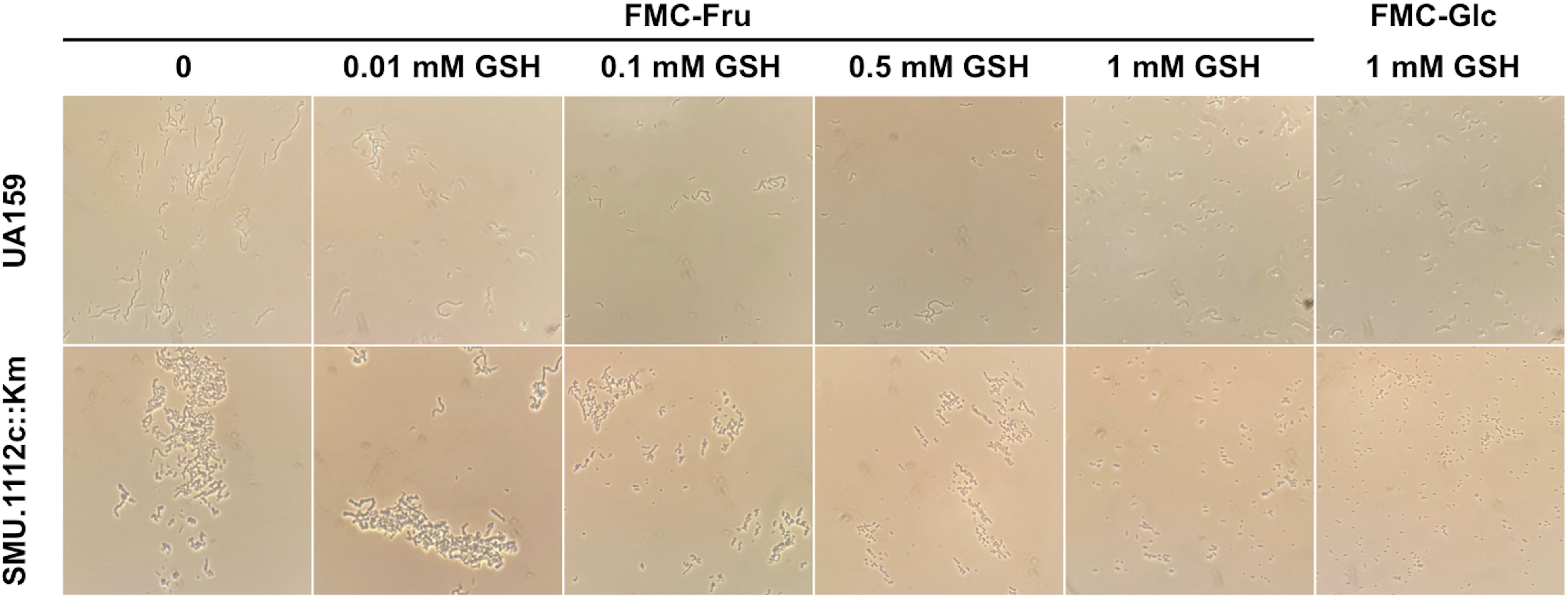
Addition of glutathione (GSH) reversed the phenotype of ΔSMU.1112c on fructose. ΔSMU.1112c was cultivated overnight with or without 1 mM of GSH, before being observed under microscope.

Interestingly, while strain ΔSMU.1112c grew very poorly on fructose (but not on mannose), strain ΔSSA_1625 showed only a minor deficiency growing on fructose, yet a more significant deficiency on mannose (Fig. S6). Further, deletion of a *fruI* orthologue in ΔSSA_1625 background failed to reverse the phenotype on mannose, although mannose (in the form of mannose-6-P) is known to be first converted to F-6-P once internalized by *S. mutans* (41). Interestingly, SSA SK36 released the most eDNA growing on mannose in comparison to glucose or fructose (Fig. S7). Further research is needed to elucidate the different roles played by these two *gloA2* orthologues in fructose- and mannose-mediated stress phenotypes in these two important oral streptococci.

Another PTS of critical importance to the pathophysiology of *S. mutans* is the glucose/mannose-PTS (EII^Man^, *manLMNO*), a major transporter for glucose and multiple other carbohydrates (42). A mutant lacking the *manL* gene (manL2020) was tested for RES tolerance. Different from mutants of the glyoxalase pathway, manL2020 showed enhanced resistance to both MG and GO when included in the MIC assay (Table 1). Additional analysis of the unique phenotype of strain manL2020 was carried out below.

### Treatment with MG significantly alters bacterial gene expression

To understand the impact of MG at the molecular level, SMU UA159 and SSA SK36 were each grown in FMC-glucose to exponential phase and treated with sub-MIC levels of MG for 30 min before harvesting. RT-qPCR assays were carried out in these cells to measure the mRNA levels of genes of the glyoxalase pathway and those deemed critical to persistence. As shown in Table 2, treatment of UA159 with MG resulted in increased expression of the putative SMU.1602-1603 operon (>2-fold), superoxide dismutase (*sod,* 7-fold)(43), glutathione synthetase (*gshAB,* 2-fold), and a fructose-inducible *rcrRPQ* operon (2-fold) that is important for *S. mutans* aerobic stress tolerance and competence (44, 45). The transcript levels SMU.1112c remained unchanged, while that of SMU.1323 showed a slight reduction. Most genes required for central carbon metabolism and PTS, including pyruvate formate lyase (*pfl*), pyruvate dehydrogenase complex (*pdh*), acetate kinase (*ackA*), glycogen synthase operon (*glg*), glucose-PTS operon (*man*), PTS EI (*ptsI*), and HPr (*ptsH*) were substantially down regulated by the treatment.

**Table 2.** RT-qPCR results showing the relative abundance of mRNA levels of important genes. Each strain was grown to exponential phase (OD_600_ = 0.5) and treated with MG (at 5 mM for SMU and 2 mM for SSA) for 30 min. Each strain was represented by at least 3 individual cultures and each cDNA was measured in technical duplicates. The results are each presented as average and standard deviation (in parenthesis). The statistical significance of each MG-treated strain was assessed against the same strain grown without MG; and for manL2020 grown without MG, against UA159 without MG. Asterisks denote statistical significance according to Student’s *t* test (*, *P* < 0.05; **, *P* < 0.01; ***, *P* < 0.001; ****, *P* < 0.0001).

| Genes | SMU UA159 |  | manL2020 |  | Genes | SSA SK36 |  |
| --- | --- | --- | --- | --- | --- | --- | --- |
|  | no MG | 5 mM MG | no MG | 5 mM MG |  | no MG | 2 mM MG |
| SMU.1112c | 1.00 (0.11) | 1.11 (0.13) | 0.90 (0.07) | 0.89 (0.07) | SSA_1625 | 1.00 (0.03) | 0.70 (0.10)* |
| SMU.1603 | 1.01 (0.14) | 2.73 (0.19)**** | 1.26 (0.16) | 3.01 (0.40)** | SSA_0962 | 1.00 (0.05) | 0.28 (0.02)*** |
| SMU.1602 | 1.00 (0.03) | 3.39 (0.36)** | 1.21 (0.12) | 3.45 (1.19) |  |  |  |
| SMU.1323 | 1.00 (0.06) | 0.81 (0.05)* | 0.86 (0.05)* | 0.54 (0.03)*** | SSA_0872 | 1.00 (0.02) | 0.52 (0.005)*** |
| *gshAB* | 1.00 (0.08) | 1.93 (0.18)** | 1.02 (0.04) | 1.69 (0.47) |  |  |  |
| *sod* | 1.00 (0.02) | 7.36 (0.75)** | 1.29 (0.04)** | 7.90 (2.50)* |  |  |  |
| *pfl* | 1.00 (0.06) | 0.20 (0.01)*** | 1.69 (0.11)*** | 0.46 (0.01)*** | *pfl* | 1.00 (0.06) | 0.12 (0.02)** |
| *pdhD* | 1.10 (0.63) | 0.37 (0.06) | 5.64 (0.63)*** | 3.10 (0.10)** | *pdhA* | 1.023 (0.26) | 1.88 (0.71) |
| *manN* | 1.01 (0.14) | 0.37 (0.05)** | 1.367 (0.12)* | 0.34 (0.01)*8 | *manL* | 1.001 (0.08) | 0.73 (0.63) |
| *ackA* | 1.01 (0.14) | 0.55 (0.07)* | 1.22 (0.08) | 0.96 (0.27) | *ackA* | 1.00 (0.06) | 1.43 (0.43) |
| *gtfB* | 1.00 (0.07) | 0.66 (0.05)** | 0.79 (0.11) | 0.33 (0.07)** | *spxB* | 1.00 (0.05) | 1.87 (0.85) |
| *glnQ* | 1.00 (0.11) | 0.38 (0.09)** | 1.11 (0.18) | 0.11 (0.01)** | *pflA* | 1.00 (0.08) | 0.85 (0.10) |
| *ptsH* | 1.00 (0.08) | 0.15 (0.01)** | 0.89 (0.09) | 0.1 (0.02)** | *nox* | 1.00 (0.06) | 1.90 (0.56) |
| *dnaK* | 1.01 (0.16) | 0.31 (0.05)* | 1.48 (0.27) | 0.16 (0.03)* | *ldh* | 1.02 (0.29) | 0.41 (0.08)* |
| *glgD* | 1.01 (0.16) | 0.39 (0.07)* | 1.73 (0.17)** | 0.76 (0.08)** | *pykF* | 1.05 (0.43) | 0.83 (0.18) |
| *rcrR* | 1.00 (0.03) | 1.88 (0.55) |  |  |  |  |  |
| *rcrP* | 1.00 (0.02) | 2.22 (0.31)* |  |  |  |  |  |
| *relA* | 1.00 (0.04) | 1.25 (0.17) |  |  |  |  |  |
| *relP* | 1.00 (0.02) | 0.90 (0.56) |  |  |  |  |  |
| *relQ* | 1.00 (0.04) | 0.43 (0.02)*** |  |  |  |  |  |
| *sloA* | 1.00 (0.06) | 0.25 (0.02)*** |  |  |  |  |  |
| *citB* | 1.01 (0.18) | 0.27 (0.04)* |  |  |  |  |  |

In comparison, treatment of SK36 with MG resulted in reduced expression in bioenergetic genes *pfl* and *ldh* (lactate dehydrogenase), although *pdh, spxB* (pyruvate oxidase), and *nox* (NADH oxidase) each showed a slight increase in mRNA levels (Table 2). Significantly different from SMU UA159, mRNA levels of the orthologue of *lguL* (SSA_0962) in SK36 were reduced 2-fold by the treatment with MG. Together, these results suggested that MG treatment at sub-MIC levels inhibits central metabolism that is required for energy production and anabolism in both bacteria, yet triggers different responses in terms of the glyoxalase pathway: *lguL* orthologue is induced in SMU but suppressed in SSA background. This could at least partly explain the greater tolerance to RES shown by the pathobiont than the commensal SSA. As depicted in Fig. 1, the genetic structure surrounding SMU.1603 appears significantly different from most oral streptococci.

To understand how the glucose-PTS mutant Δ*manL* of SMU UA159 presents enhanced RES resistance, the same treatment and RT-qPCR analysis were performed on SMU mutant manL2020. When treated with MG, the *manL* mutant showed (Table 2) wild-type levels of expression by SMU.1112c, *lguL*, and SMU.1602, *gshA*, *sod*, and several other pathways. However, unlike the wild type, increased expression of several metabolic pathways/genes, *glgD*, *ackA,* but chiefly the *pdh* operon, was observed, revealing superior energy management under RES stress and providing a plausible explanation for enhanced RES tolerance by the *manL* mutant. Previous research using *S. mutans* suggested that an intact PDH complex is essential to bacterial survival under stress such as extended starvation (46) and acidic conditions (47). Notably, similarly constructed glucose-PTS (*manL*) mutants in the backgrounds of SSA SK36 and SGO DL1 each showed a slight reduction in MG tolerance (Table 1 and Fig. S3).

### MG metabolism influences interspecies competition

An important objective of this study was to assess the effect of RES on microbial interactions between *S. mutans* and commensals. A growth competition between SMU UA159 and SSA SK36, each labeled with a different antibiotic marker, was carried out in a planktonic culture prepared using FMC supplemented with glucose or fructose, with or without addition of MG. As indicated in Fig. 8, when co-cultured without MG, both species appeared comparable in competitiveness, either in glucose-or fructose-based medium. However, when 1 mM MG was added to the media, SMU UA159 showed 20∼170-fold increases in competition indices depending on the sugar, a result consistent with our earlier finding that SMU UA159 is significantly more resistant to MG than SSA SK36.

**Fig. 8.**
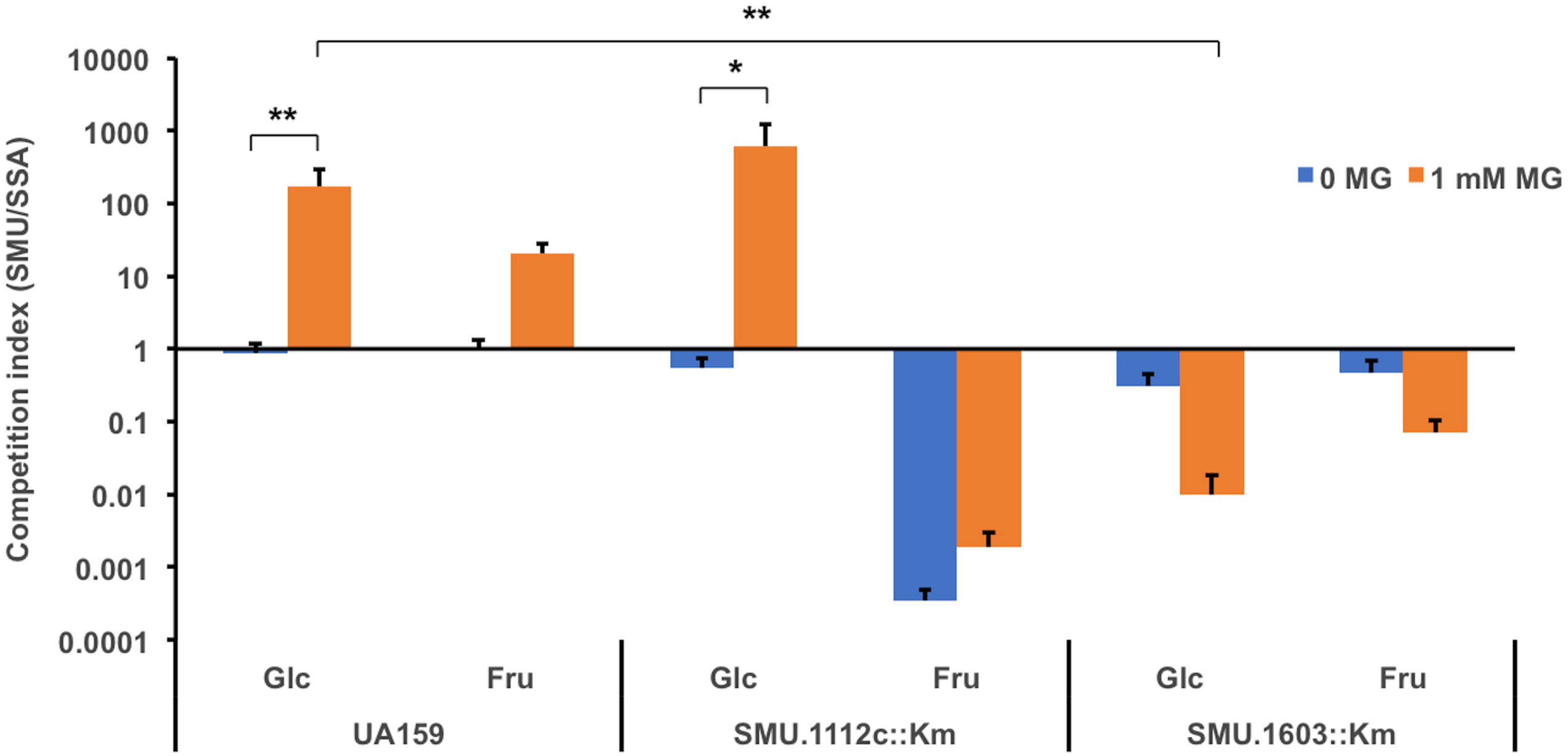
Competition between *S. mutans* and *S. sanguinis*. Exponential-phase cultures of SMU UA159-Km and ΔSMU.1112c and ΔSMU.1603 were each mixed with SSA MMZ1945 at 1:1 ratio, followed by dilution at 1000-fold into FMC containing 20 mM or glucose or fructose, and incubation in 5% CO_2_ atmosphere for 24 h. CFU enumeration at both the start and the end of the incubation resulted in competition indices of each species. Each strain was represented by at least three independent cultures, and the experiment was repeated twice. Statistical significance was assessed using three-way ANOVA (*, *P* < 0.05; **, *P* < 0.01).

We then tested the role of SMU.1112c and SMU.1603 in this competition by replacing the wild-type UA159 with their respective mutants. As shown in Fig. 8, loss of SMU.1112c resulted in little change to the relative competitiveness between these two species in FMC-glucose, regardless the presence of MG. However, in FMC-fructose, strain ΔSMU.1112c was markedly less competitive relative to the wild type, showing a ∼3000-fold reduction in competition indices without MG, but an even greater reduction (∼11,000-fold) when treated with 1 mM MG. This outcome echoed our earlier conclusion that SMU.1112c is primarily responsible for fitness when fructose is the primary growth carbohydrate, but may also deal with RES stress. Conversely, when strain ΔSMU.1603 was used to substitute SMU UA159, little difference was observed in its competitiveness against SSA SK36 in the absence of MG. When presented with 1 mM MG, ΔSMU.1603 showed about ∼3000-fold reduction in competitiveness in medium containing fructose, but its competition index was ∼17,000-fold lower than the wild type on glucose.

### Concluding remarks

We reported here a systematic characterization of streptococcal glyoxalase pathway that is essential to methylglyoxal and glyoxal tolerance in the oral microbiome, where RES compounds are constantly being produced by microbes and their host, especially during glycolysis. In response to our sugar-rich Western diet and/or diabetic physiology, certain constituents of the oral microbiota could have evolved novel genetics that affords them advantages during competition with other organisms by becoming better at tolerating or detoxifying RES. Results of our study suggested that *S. mutans* could outcompete commensal streptococci in the presence of RES by several mechanisms. First, the genetic structure in SMU where *lguL/*SMU.1603 finds itself includes an uncharacterized redox-modulating enzyme, a putative efflux pump protein, and a likely transcription regulator, all of which could aid in the expression and functionality of LguL in not-yet-understood ways. This genetic structure is present in a few other streptococcal species, but absent in most commensals analyzed. The fact that the *lguL* mutants in SMU and SSA backgrounds showed comparable tolerance to MG strongly suggests that the SMU.1602-1605 locus and its regulation is critical to the apparently greater RES resistance of *S. mutans*. Second, although not yet fully understood, SMU.1112c contributes to fructose and MG tolerance of SMU in a manner related to cellular GSH::GSSG balance. It is understood in mammalian cells that fructose oxidation in the presence of H_2_O_2_ worsens RES-mediated cellular injury (38, 48). It is thus probable that peroxigenic commensals such as SSA and SGO may be uniquely vulnerable to RES damage due to higher levels of cellular H_2_O_2_, especially when presented with fructose. Conversely, Gram-negative bacterium *E. coli* is known to activate a proton importer for cytoplasmic acidification as a means of blunting MG-induced cellular damage, in part by improving protonation of proteins or other cellular macromolecules (12). Such a mechanism is likely absent in streptococci based on genomic analysis, raising the possibility that lactic acid bacteria as avid producers of organic acids might be innately more tolerant to RES effects. If so, cariogenic, aciduric pathobionts such as *S. mutans* and lactobacilli could be naturally even better protected from RES. Last, the glucose-PTS (EII^Man^) of both *S. mutans* and *S. sanguinis* have been shown to regulate cellular bioenergetics and competitiveness, with significant distinctions in both underlying mechanism and phenotype (49, 50). The distinct effects on MG tolerance caused by *manL* deletion in these streptococci suggest different roles of PTS in RES-related gene regulation. In conclusion, RES as an important group of metabolic byproducts are severely understudied in the context of oral microbiology, considering the etiology of dental caries and the epidemic of hyperglycemic disorders affecting the populations of the world. RES are likely more abundant when conditions favor caries development, thus the competitive advantage of *S. mutans* and other cariogenic pathobionts, if proven, in the presence of RES may be a major contributor to the ecological shifts that are characteristic of the development of cariogenic microbiomes. Future research into the contributions by, and mechanisms involved in regulating, RES-detoxifying pathways in pathobionts as well as oral microbiome could provide important knowledge for oral health and better caries prevention, especially for diabetic individuals.

## Materials and Methods

### Bacterial strains and culture conditions

*S. mutans* UA159, *S. sanguinis* SK36, *S. gordonii* DL1, their genetic derivatives (Table 3), and various other wild-type oral streptococcal isolates were first refreshed from frozen stocks by growing on BHI (Difco Laboratories, Detroit, MI) agar plates, then overnight in liquid BHI medium, before being diluted into fresh BHI, Tryptone-yeast extract medium (TY, 30 g of Tryptone and 5 g of yeast extract per liter), or FMC synthetic medium (51) that was supported with various carbohydrates as specified by each assay. Antibiotics were used, when necessary, at the following concentrations: kanamycin (Km) 0.5 to 1 mg/ml; erythromycin (Em) 5 to 10 µg/ml. All agar plates and liquid cultures were incubated at 37°C in an ambient atmosphere maintained with 5% CO_2_ unless specified otherwise. Bacterial morphology in liquid cultures was assessed, without staining, using a Nikon Eclipse E400 microscope under the phase-contrast setting. For analysis of bacterial growth, BHI cultures of individual strains from the exponential phase were diluted 100-fold into FMC media in a 96-well plate, each at 200 µl volume and covered with 60 µl of mineral oil, before being loaded onto a Synergy 2 plate reader (BioTek, Agilent Technologies, Santa Clara, CA) maintained at 37°C, with optical density (λ = 600 nm, OD_600_) data collected at 1-h interval. Antibiotics, sodium glutathione, methylglyoxal (MG), and glyoxal (GO) were purchased from MilliporeSigma (Burlington, MA).

**Table 3.** Strains used in this study (excluding oral commensal isolates in Table 1).

| Strains | Relevant characteristics<sup>a</sup> | Source or reference |
| --- | --- | --- |
| UA159 | *S. mutans* wild type, *perR*<sup>+</sup> | ATCC 700610 |
| manL2020 | UA159 *manL::Km* | From UA159 |
| ΔSMU.1112c::*Km* | UA159 SMU.1112c::*Km* | From UA159 |
| ΔSMU.1112c::*Em* | UA159 SMU.1112c::*Em* | From UA159 |
| ΔSMU.1112cCom | ΔSMU.1112c complementation | From ΔSMU.1112c::*Km* |
| ΔSMU.1323 | UA159 SMU.1323::*Km* | From UA159 |
| ΔSMU.1323Com | ΔSMU.1323 complementation | From ΔSMU.1323::*Km* |
| ΔSMU.1603 | UA159 SMU.1603::*Km* | From UA159 |
| ΔSMU.1603Com | ΔSMU.16033 complementation | From ΔSMU.1603::*Km* |
| ΔSMU.1112c/*frul* | UA159 SMU.1112c::*Km frul::Em* | From ΔSMU.1112c::*Km* |
| ΔSMU.1112c/*levD* | UA159 SMU.1112c::*Km levD::Em* | From ΔSMU.1112c::*Km* |
| ΔSMU.1112c/1603 | UA159 SMU.1112c::*Km* SMU.1603::*Em* | From ΔSMU.1112c::*Km* |
| ΔSMU.1323/1603 | UA159 SMU.1323::*Km* SMU.1603::*Em* | From ΔSMU.1323::*Km* |
| Δ*gshAB* | UA159 *gshAB::Km* | From UA159 |
| Δ*gshAB*Com | Δ*gshAB* complementation | From Δ*gshAB* |
| UA159-Km | *gtfA::Km* | (53) |
| SK36 | *S. sanguinis* wild type | Kitten laboratory |
| manL1617 | SK36 *manL::Km* | (50) |
| MMZ1945 | SK36 *gtfP::Em* | From SK36 |
| ΔSSA_0962 | SK36 SSA_0962::*Km* | From SK36 |
| ΔSSA_1625 | SK36 SSA_1625::*Km* | From SK36 |
| DL1 | *S. gordonii* wild type | ATCC 49818 |
| manL898 | DL1 *manL::Km* | DL1 (60) |

746 *Em*, and *Km*: resistance against erythromycin and kanamycin, respectively.

### Construction of genetic mutants and complementation

Deletion of genes of interest was conducted by allelic exchange strategy using a mutator DNA that was assembled (Gibson assembly) from two flanking sequences and an antibiotic marker in between, which would replace the target gene with the antibiotic cassette through homologous recombination (52). Transformation of the wild type or mutant strains was achieved using naturally competent cells induced by treatment with competence-stimulating peptide (CSP) for respective species when growing in BHI. To improve scientific rigor, we routinely utilized a nonpolar kanamycin marker (*Km*), which lacked a promoter sequence, and an erythromycin marker (*Em*) with its own promoter, to knock out the same gene separately; both mutants were assayed for phenotypic change.

Complementation analysis for phenotypic validation in mutants was carried out by a knock-in strategy that replaced in the mutator DNA the initial antibiotic cassette with an alternative marker, along with a wild-type copy of the target gene in between the two flanking fragments, followed by transformation of the said mutant. PCR amplification and assembly of the mutator DNA via Gibson assembly was performed according to a previously published protocol detailed elsewhere (50). All genetic derivatives constructed in this study have been confirmed by PCR reactions targeting regions of interest, followed by Sanger sequencing. All oligonucleotides used in this procedure are listed in Table S1.

### Minimum inhibitory concentration (MIC) assay

To measure the MIC for MG or GO, bacterial strains were first cultured overnight in BHI medium supplemented with 10 mM of potassium phosphate buffer (pH 7.2), then diluted 40-fold into 200 µl of FMC medium supported with 20 mM of glucose and varying concentrations of MG or GO. After incubating in an ambient incubator maintained with 5% of CO_2_ at 37°C for 20 to 24 h, the optical density (OD_600_) of the cultures were measured using a plate reader. Each strain was represented by at least three individual cultures. The minimum inhibitory concentration was defined as the levels above which OD_600_ remained unchanged relative to the blank. To report the increment in concentration used in the assay, the average MIC value is presented as a range, with the second value denoting the next immediate concentration above the actual MIC.

### Planktonic growth competition assay

To evaluate bacterial competitiveness when cultivated together in a planktonic setting, a WT SMU strain, UA159-Km (53) or other deletion mutants (Table 3), was marked with a Km marker while an SSA strain MMZ1945 was marked by an Em marker at the *gtfP* site (54). Each strain was grown overnight in BHI (n = 3), followed by sub-culturing in the same BHI medium till exponential phase (OD_600_ = 0.5). After mixing SMU and SSA strains at 1:1 volume, the cells were diluted 1000-fold into FMC medium supplemented with 20 mM of glucose or fructose, in addition to 0 or 1 mM of MG. The diluted cultures were then incubated for 24 h in 95% air and 5% CO_2_ at 37°C. Both before (T_0_) and after the 24-h incubation (T_24_), the mixed cultures were subjected to a 15-second sonication at 100% power (FB120 water bath sonicator, Fisher Scientific), followed by serial dilution and plating onto selective agar plates containing either Km or Em. All plates were incubated for 2 days before CFU enumeration and calculation of competition indices. The competition index (SMU over SSA) was calculated as [SMU(T_24_)/ SSA(T_24_)] / [SMU(T_0_)/ SSA(T_0_)], with values >1 indicating SMU being more competitive than SSA, and vice versa.

### Biofilm assay

Development of biofilms was conducted on abiotic surfaces following previously published protocols (55). In brief, bacterial cultures from mid-exponential phase (OD_600_ = 0.5) were diluted 100-fold into a biofilm medium BM (56) supplemented with 2 mM sucrose and 18 mM glucose (BMGS) or fructose (BMFS). The culture dilutions were then loaded, at 200 μl/well, onto a 96-well polystyrene microplate (Corning 3917) and incubated in 95% air and 5% CO_2_ for 2 days without agitation. Biofilms were then stained with crystal violet to assess the total biomass, which was quantified by elution of the stain with 30% acetic acid and measurement of its optical density at 575 nm.

### RNA extraction and RT-qPCR

RNA extraction and mRNA quantification was conducted by following a previously described protocol (57). Briefly, bacterial cultures were prepared as above to exponential phase (OD_600_ = 0.4) in FMC medium containing 20 mM glucose, added 0 or 1 mM of MG and returned to incubation for 30 min, before being harvested by centrifugation. After treatment with RNAprotect Bacteria reagent, bacterial cell envelope was disrupted by rapid homogenization in the presence of glass beads, SDS, and acidic phenol and chloroform, followed by centrifugation to separate cell debris. Clarified cell lysate was processed using a Qiagen RNeasy kit and in-column treatment with a DNase I kit (Qiagen, Germantown, MD) for purification of total RNA. For RT-qPCR, a reverse transcription kit (iScript select cDNA synthesis kit, Bio-Rad, Hercules, CA) was used to create cDNA from the total RNA with gene-specific antisense primers (Table S1), followed by Real-time PCR analysis using the CFX96 system and SYBR Green Supermix (Bio-Rad). Relative mRNA levels of each gene were quantified using the ΔΔCq method and an internal reference gene *gyrA*.

### Statistical analysis and data availability

Statistical analysis of data was carried out using the software of Prism (GraphPad of Dotmatics, San Diego, CA). Any data, strains, and materials generated by this study will be available upon request from the authors for research or validation.

## Acknowledgements

This study was supported by a grant from NIDCR to Lin Zeng and Robert A. Burne (DE012236) and a startup fund from UF Office of Research to Lin Zeng. We thank Alexandra Peterson and Zachary Taylor for technical assistance in generation of some of the graphs used in this study.

## Author Contributions

LZ designed the study; LZ, PN, and BG performed the experiments; LZ and PN analyzed the data; LZ and RAB wrote the manuscript.

